# Predicting walking response to ankle exoskeletons using data-driven models

**DOI:** 10.1101/2020.06.18.105163

**Authors:** Michael C. Rosenberg, Bora S. Banjanin, Samuel A. Burden, Katherine M. Steele

## Abstract

Despite recent innovations in exoskeleton design and control, predicting subject-specific impacts of exoskeletons on gait remains challenging. We evaluated the ability of three classes of subject-specific phase-varying models to predict kinematic and myoelectric responses to ankle exoskeletons during walking, without requiring prior knowledge of specific user characteristics. Each model – phase-varying (PV), linear phase-varying (LPV), and nonlinear phase-varying (NPV) – leveraged Floquet Theory to predict deviations from a nominal gait cycle due to exoskeleton torque, though the models differed in complexity and expected prediction accuracy. For twelve unimpaired adults walking with bilateral passive ankle exoskeletons, we predicted kinematics and muscle activity in response to three exoskeleton torque conditions. The LPV model’s predictions were more accurate than the PV model when predicting less than 12.5% of a stride in the future and explained 49–70% of the variance in hip, knee, and ankle kinematic responses to torque. The LPV model also predicted kinematic responses with similar accuracy to the more-complex NPV model. Myoelectric responses were challenging to predict with all models, explaining at most 10% of the variance in responses. This work highlights the potential of data-driven phase-varying models to predict complex subject-specific responses to ankle exoskeletons and inform device design and control.

## III. Introduction

Ankle exoskeletons are used to improve kinematics and reduce the energetic demands of locomotion in unimpaired adults and individuals with neurologic injuries [1–5]. Customizing exoskeleton properties to improve an individual’s gait is challenging and accelerating the iterative experimental process of device optimization is an active area of research [6, 7]. Studies examining the effects of exoskeleton properties – sagittal-plane ankle stiffness or equilibrium ankle angle for passive exoskeletons and torque control laws for powered exoskeletons – on kinematics, motor control, and energetics have developed design and control principles to reduce the energetic demand of walking and improve the quality of gait [1, 6, 8, 9]. Predicting how an individual’s gait pattern responds to ankle exoskeletons across stance may inform exoskeleton design by enabling rapid evaluation of exoskeleton properties not tested experimentally. Additionally for powered exoskeletons, which prescribe torque profiles using feedforward or feedback (e.g. kinematic or myoelectric) control laws, predicting responses over even 10–20% of a stride may improve tracking performance or transitions between control modes [4, 10–12]. However, predicting subject-specific responses to exoskeletons remains challenging for unimpaired individuals and those with motor impairments [2, 12, 13].

Common physics-based models, including simple mechanical models and more physiologically-detailed musculoskeletal models, use principles from physics and biology to analyze and predict exoskeleton impacts on gait. For example, one lower-limb mechanical walking model predicted that an intermediate stiffness in a passive exoskeleton would minimize the energy required to walk, a finding that was later observed experimentally in unimpaired adults [1, 14]. More physiologically-detailed musculoskeletal models have been used to predict the impacts of exoskeleton design on muscle activity during walking in children with cerebral palsy and running in unimpaired adults [15, 16]. While these studies identified hypothetical relationships between kinematics and the myoelectric impacts of exoskeleton design parameters, their predictions were not evaluated against experimental data.

Challenges to accurately predicting responses to ankle exoskeletons with physics-based models largely stem from uncertainty in adaptation, musculoskeletal physiology, and motor control, which vary between individuals and influence response to exoskeletons. While individuals explore different gait patterns to identify an energetically-optimal gait, exploration does not always occur spontaneously, resulting in sub-optimal gait patterns for some users [17]. Popular physiologically-detailed models of human gait typically assume instantaneous and optimal adaptation, which do not reflect how experience and exploration may influence responses to exoskeletons, possibly reducing the accuracy of predicted responses [18, 19]. Additionally, when specific measurement sets are unavailable for model parameter tuning, population-average based assumptions about musculoskeletal properties and motor control are required [17, 20–22]. However, musculoskeletal properties and motor control are highly uncertain for individuals with motor impairments, today’s most ubiquitous ankle exoskeleton users [19, 20, 22, 23]. Musculotendon dynamics and motor complexity are known to explain unintuitive exoskeleton impacts on gait energetics, suggesting that uncertain musculotendon parameters and motor control may limit the accuracy of predicted changes in gait with ankle exoskeletons [19, 21, 24]. Predictions of exoskeleton impacts on gait using physiological models, therefore, require accurate estimates of adaptation, musculotendon parameters, and motor control.

Conversely, data-driven approaches address uncertainty in user-exoskeleton dynamics by representing the system entirely from experimental data. For instance, human-in-the-loop optimization provides a model-free alternative to physics-based prediction of exoskeleton responses by automatically exploring different exoskeleton torque control strategies for an individual [6, 7]. This experimental approach requires no prior knowledge about the individual: optimization frameworks identify torque control laws that decrease metabolic rate relative to baseline for an individual using only respiratory data and exoskeleton torque measurements. However, experimental approaches to exoskeleton optimization require the optimal design to be tested, potentially making the search for optimal device parameters time-intensive. Alternatively, machine learning algorithms, such as the Random Forest Algorithm, have used retrospective gait analysis and clinical exam data to predict changes in joint kinematics in response to different ankle-foot orthosis designs in children with cerebral palsy [8]. This study reported good classification accuracy, though predictions may not generalize to new orthosis designs. Unlike physiologically-detailed or physics-based models, human-in-the-loop optimization and many machine learning models are challenging to interpret, limiting insight into how a specific individual’s physiology influences response to exoskeleton torque. A balance between physiologically-detailed and model-free or black-box data-driven approaches may facilitate the prediction and analysis of responses to ankle exoskeletons without requiring extensive knowledge of an individual’s physiology.

In this work, we investigated a subject-specific data-driven modeling framework – phase-varying models – that may fill the gap between physiologically-detailed model-based and model-free experimental approaches for predicting gait with exoskeletons. Phase-varying models typically have linear structure whose parameters are estimated from data, enabling both prediction and analysis of gait with exoskeletons [25, 26]. Unlike physiologically-detailed models, phase-varying models do not require knowledge of the physics or control of the underlying system. Unlike experimental approaches, the model-based framework enables prediction of responses to untested exoskeleton designs or control laws.

Phase-varying models leverage dynamical properties of stable gaits derived from Floquet Theory, which ensures that the convergence of a perturbed trajectory to a stable limit cycle may be locally approximated using time-varying linear maps [27]. Similar principles have been shown to generalize to limit cycles in non-smooth or hybrid systems, such as human walking [28]. Moreover, phase-varying modeling principles have been applied to biological systems, identifying linear phase-varying dynamics to investigate gait stability and predict changes in kinematics in response to perturbations [25, 26, 29–31]. Responses to ankle exoskeleton torques may be similarly defined as perturbations off an unperturbed (*i.e.* zero torque) gait cycle, suggesting that the principles of phase-varying models will generalize to walking with exoskeletons. To the best of our knowledge, phase-varying models have never been used to study walking with exoskeletons and the extent to which the principles underlying phase-varying models of locomotion generalize to walking with exoskeletons is unknown.

To determine if phase-varying models represent useful predictive tools for locomotion with exoskeletons, the purpose of this research was to evaluate the ability of subject-specific phase-varying models to predict kinematic and myoelectric responses to ankle exoskeleton torque during walking. We predicted responses to exoskeletons in unimpaired adults walking with passive ankle exoskeletons under multiple dorsiflexion stiffness conditions. We focused on three related classes of phase-varying models with different structures, complexity, and expected prediction accuracies: a phase-varying (PV), a linear phase-varying (LPV), and a nonlinear phase-varying (NPV) model. Since passive exoskeletons typically elicit small changes in joint kinematics and muscle activity, we expected the validity of Floquet Theory for human gait to extend to gait with exoskeletons, indicating that the LPV model should accurately predict responses to passive exoskeleton torque [1, 25–27, 29]. We, therefore, hypothesized that the LPV models would predict kinematic and myoelectric responses to torque more accurately than the PV model and as accurately as the NPV model. To exemplify the potential utility of subject-specific phase-varying models in gait analysis with ankle exoskeletons, we show how varying the length of model prediction time horizon may inform measurement selection for exoskeleton design and control. To assess the viability of data-driven phase-varying models in gait analysis settings, we evaluated the effect of limiting the size of the training dataset on prediction accuracy.

## IV. Methods

### A. Experimental protocol

We collected kinematic and electromyographic (EMG) data from 12 unimpaired adults (6 female / 6 male; age = 23.9 ± 1.8 years; height = 1.69 ± 0.10 m; mass = 66.5 ± 11.7 kg) during treadmill walking with bilateral passive ankle exoskeletons at a self-selected speed. Each participant performed two sessions on separate days within a one week span. In the first session, we modified the exoskeletons for fit and comfort and performed a 20-minute practice session. *Additional detail regarding experimental setup, input variable calculations, modeling algorithms, and statistical analyses can be found in Supplemental – S1*.

Data were collected during the second session. We monitored changes in kinematics using a modified Helen-Hayes marker set [32] and a 10-camera motion capture system (Qualisys AB, Gothenburg, SE), and measured muscle activity using 14 wireless EMG sensors (Delsys Inc., Natick, USA). The EMG sensors were placed bilaterally on the soleus, medial gastrocnemius, tibialis anterior, vastus medialis, rectus femoris, lateral hamstrings, and gluteus medius following SENIAM guidelines [33]. Participants performed four randomized trials on a split-belt instrumented treadmill (Bertec Corp., Columbus, USA) under different exoskeleton conditions (Fig. 1). Unlike many clinical exoskeletons (ankle-foot orthoses), whose torque profiles are smooth functions of ankle angle, the passive exoskeletons used in this study generated ankle plantarflexion torques as a piecewise-linear function of the user’s ankle angle, and the exoskeleton’s neutral angle and rotational stiffness. The exoskeletons did not resist plantarflexion, similar to other experimental devices [1, 3]. The four exoskeleton conditions were set to sagittal-plane stiffness values: K_0_ (0 Nm/deg), K_1_ (1.17 Nm/deg), K_2_ (3.26 Nm/deg), and K_3_ (5.08 Nm/deg), a range known to alter kinematics and myoelectric signals during gait (Fig. 1) [1]. Participants walked for six minutes per trial, the last four of which were recorded, and could rest between trials.

**Fig. 1.**
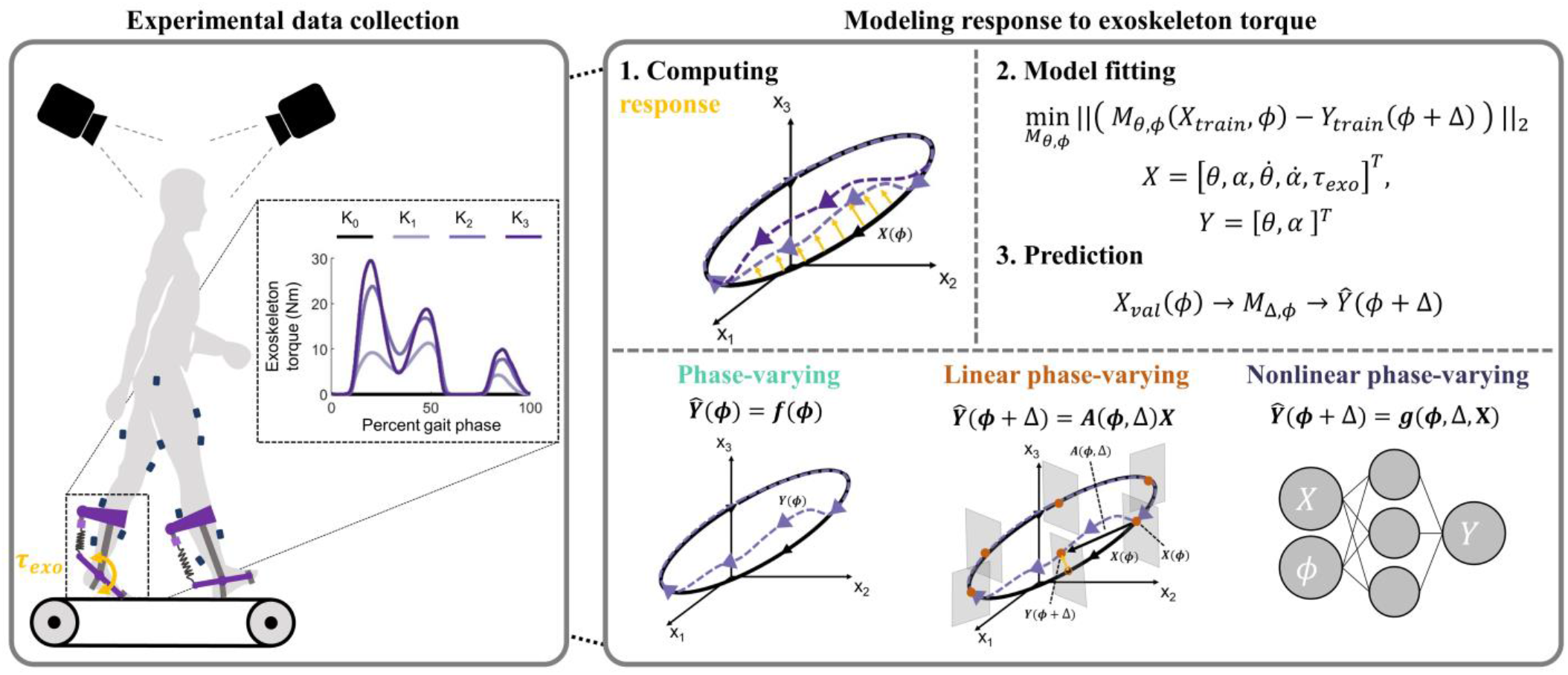
<u>Left box</u>: Data were collected during treadmill walking with bilateral ankle exoskeletons that used linear springs to resist dorsiflexion. Increasing exoskeleton stiffness (K_0_–K_3_) increased exoskeleton torque (τ_exo_, yellow). <u>Right box</u>: (1) Purple dashed arrows represent responses to exoskeleton torque, which were defined as deviations from the average zero-torque gait cycle (K_0_). (2) Response data from the training set were used to fit each model. Input variables included joint kinematics, muscle activity, their time derivatives, and exoskeleton torque. (3): Models were validated by predicting responses from the held-out torque condition using the models fit in (2). <u>Right box (bottom)</u>: The three phase-varying models were fit and evaluated on the same training and validation sets. *M*_*Δϕ*_ = generic model function of prediction horizon and phase; X = experimental inputs; Y = experimental outputs; 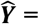 predicted outputs; *ϕ* = phase; Δ = prediction horizon; A = linear function; f, g = nonlinear functions; Δ = joint kinematics; *α* = muscle activation; τ_*exo*_ = exoskeleton torque.

The marker trajectories were low-pass filtered at 6 Hz using a zero-lag fourth-order Butterworth filter [5]. We computed joint kinematics by scaling a generic 29 degree-of-freedom skeletal model to each participant’s skeletal geometry and body mass using the inverse kinematics algorithm in OpenSim 3.3 to convert marker trajectories into joint kinematics [18, 34]. To compute linear EMG envelopes, we high-pass filtered the EMG data at 40 Hz, rectified the data, and low-pass filtered at 10 Hz [9]. Kinematic and EMG data were pre-processed using custom scripts in MATLAB (MathWorks, Natick, USA).

### B. Gait phase and phase-varying models

Unlike the typical gait cycle definition – the percentage of time between successive foot contact events – we used a gait phase based on kinematic posture, which we expected to improve predictions of a system’s response to perturbations from the exoskeletons [35]. Using a posture-based gait phase groups kinematically-similar measurements at a specific phase, reducing variance in the data at any point in the cycle, and ensuring that similar postures across exoskeleton conditions were used during model fitting and prediction. Moreover, Floquet Theory ensures that phase is well-defined using any periodically-varying measurements [27]. We used the *Phaser* algorithm, which estimates a system’s phase using arbitrary input signals considered to be phase-locked, to generate gait phase estimates as a function of left and right leg hip flexion angles, similar to a phase variable proposed to control robotic prostheses [30, 35]. Following gait phase estimation, we modeled gait using three subject-specific models of response to exoskeletons:

#### 1) Phase-varying model

The phase-varying (PV) model was our simplest model and predicts outputs purely as a function of gait phase. Rather than taking exoskeleton torque as an input, PV model predictions are similar to guessing the average of the training data at a certain gait phase (Table I) [30, 36]. The PV model takes a phase estimate as an input and returns a prediction of the system’s outputs, 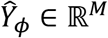, where *M* is the number of outputs. The PV model was parameterized using a seventh-order Fourier Series as a function of phase and served as a lower bound on prediction accuracy.

**Table I:** Summary of model structures and expected prediction accuracies.

| Model | Functional form | Linear terms | Nonlinear terms | Expected prediction accuracy |
| --- | --- | --- | --- | --- |
| Phase-varying (PV) | $\hat{Y}_\phi = F_{PV}(\phi)$ | None | Phase | Low |
| Linear phase-varying (LPV) | $\hat{Y}_{\phi+\Delta} = F_{LPV,\Delta}(\phi)X_\phi$ | Inputs | Phase | Moderate |
| Nonlinear phase-varying (NPV) | $\hat{Y}_{\phi+\Delta} = G_{NPV,\Delta}(\phi, X_\phi)$ | None | Phase<br>Inputs | Moderate-High |

$F$ = model functions parameterized by a Fourier Series; $G$ = feedforward neural network model; $\phi$ = phase; $\Delta$ = prediction horizon; $X$ = inputs; $\hat{Y}$ = predicted outputs

#### 2) Linear phase-varying model

The linear phase-varying (LPV) model is a discrete-time model that predicts system outputs at a future phase based on measurements at an initial phase (Table I). For any phase, *ϕ*, from 0-100% of a stride and a prediction horizon, *Δ*, the LPV model estimates a map *A*_*ϕ,Δ*_ ∈ ℝ^*M×N*+1^, from the initial phase to the final phase, where *N* + 1 denotes the number of input variables (*N*) plus a constant term. At 64 initial phases spaced equally over the gait cycle, we fit discrete maps between initial and final phases using weighted least-squares regression [25, 26, 29]. We weighted each observation based on the proximity of its phase estimate to the prescribed initial phase using a Gaussian weighting scheme. For each prediction horizon, the LPV model was represented as a continuously phase-varying function, *F*_*LPV,Δ*_(*ϕ*) ≈ *A*_*ϕ,Δ*_, parametrized by a Fourier Series. We expected the LPV model’s prediction accuracy to exceed that of the PV model [27].

### 3) Nonlinear phase-varying model

While the LPV model should approximate nonlinearities in the dynamics of response to torque, we selected a nonlinear phase-varying (NPV) model that serves as an upper bound on prediction accuracy. Specifically, we used a three-layer feedforward neural network – a universal function approximator (Table I) [37]. Neural networks are considered state-of-the-art predictors and are used in numerous domains, including image recognition and robotics [38]. The NPV model’s parameters were tuned for each prediction horizon and included phase as an input. We expected the NPV model’s prediction accuracy to meet or exceed that of the LPV model.

### C. Inputs and output variables

To reflect clinically-relevant measurements and the dynamics of the neuromusculoskeletal system, we selected input variables expected to encode musculoskeletal dynamics and motor control: 3D pelvis orientation and lower-limb and lumbar joint angles, processed EMG signals, and their time derivatives at an initial phase, *ϕ* [1, 2, 39]. We appended ten time-history exoskeleton torque samples per leg – uniformly distributed between the initial and final phases – to the inputs, resulting in *N* = 80 inputs [6, 12]. Our decision to use exoskeleton torque samples was motivated by Floquet Theory, according to which an individual’s posture at a future time is a linear function of their initial posture and the exoskeleton torque signal between initial and final times [27]. Model outputs (*M* = 20) included right and left leg sagittal-plane hip, knee and ankle kinematics, and EMG signals from each muscle at a future phase, *ϕ* + *Δ*, offset from the initial phase by prediction horizon *Δ*. While phase-varying models may also predict joint moments, we omitted prediction of kinetic outcomes due to the presence of sporadic poor force plate strikes for some gait cycles in our dataset. We modeled response to exoskeleton torque as the deviation from the unperturbed gait cycle (*i.e.* the zero-torque, K_0_ condition) by subtracting the phase-averaged zero-torque gait cycle from each exoskeleton condition [26, 29]. All data were de-meaned and scaled to unit variance of the training set. *Additional detail regarding the selection of torque as model inputs and experimental ground reaction forces can be found in Supplemental – S2.*

We first computed each model’s ability to predict responses to torque within the range of exoskeleton stiffness levels used to train the models (interpolation) by training each model using the K_0_, K_1_ and K_3_ datasets and validating by predicting outputs from the held-out K_2_ dataset using inputs from the same dataset at an initial gait phase. While “what-if” predictions – predicting responses to “untested” (held-out) torques using nominal kinematics and EMG from a “tested” condition (e.g. K_0_) – are needed to for predictions to inform passive exoskeleton design, we chose to predict using the held-out inputs to provide unambiguous interpretation of each model’s prediction accuracy. In “what-if” predictions, errors stem from both poor model fit and poor matches between the “tested” and “untested” input data at the initial phase. By instead predicting using “untested” inputs, our predictions errors reflect only the models’ fits to each participant’s dynamics and provide upper bounds on the potential accuracy of “what-if” predictions. We selected the K_2_ condition for validation in our experimental design because responses in this intermediate torque condition should be encoded by the K_0_, K_1_, and K_3_ conditions. During validation, experimental outputs from the K_2_ condition were compared to the corresponding model predictions.

We quantified each model’s prediction accuracy using the relative remaining variance (RRV) of model predictions compared to the held-out experimental data [25]. The RRV is calculated as the ratio of the variances of the prediction error and the experimental data. An RRV value of zero implies a perfect prediction, while unity RRV values can be achieved by predicting the mean of the validation data. Since we de-meaned the data and predicted deviations from the zero-torque condition, RRV values below unity indicate that predictions are more accurate than guessing constant (e.g. zero) response to exoskeleton torque. We computed RRV values for each output using a bootstrapping procedure with 200 iterations [25]. We computed RRV values for each model over the entire validation time series of approximately 240 strides. During analysis, the right and left leg RRV values for each output variable were averaged, as we expected nearly symmetric responses from our unimpaired participants.

We evaluated the LPV and NPV models’ prediction accuracies for the K_2_ condition over a range of prediction horizons, in increments of 6.25% (1/16^th^ of a stride), between 6.25 and 100% of a gait cycle. When optimizing exoskeleton torque profiles, predicting responses using measurements at an initial phase (e.g. initial contact of the foot with the ground) to achieve a desired outcome at some final phase may be of interest, such as improving midstance knee kinematics in children with cerebral palsy [2, 3]. However, as the prediction horizon increases, coherence between measurements at initial and final phases decreases due to nonlinearities in musculoskeletal dynamics, resulting in prediction accuracies reducing to those of the average prediction (*i.e.* the PV model), rather than a stride-specific prediction [39–41]. Identifying the maximum prediction horizon at which initial measurements improve predictions at a final phase may inform exoskeleton control laws or design criteria. Therefore, we identified the largest prediction horizon lengths at which RRV values were significantly less than those of the PV model, which were constant across prediction horizons.

The amount of data required to accurately predict response to exoskeletons will restrict the settings in which phase-varying models are practical, such as in clinical gait analysis where datasets typically contain only a few gait cycles [2, 8]. We quantified the impact of training set size on prediction accuracy by determining the amount of training data needed for prediction accuracies of the K_2_ condition to approach to their values when models were fit using the entire training set (*RRV*_*full*_). We iteratively reduced the training set size by 10% of the full size (approximately 24 strides per exoskeleton condition), removing data from the end of each torque condition in the training set, providing a range of 24-240 strides of training data per condition. For all training set sizes, we evaluated models using the full-length validation set.

To test each model’s generalizability across a range of exoskeleton torque conditions, we separately predicted responses to torque in the K_1_, K_2_, and K_3_ datasets, termed *held-out conditions*, at a 12.5% stride prediction horizon (1/8^th^ of a stride). Predictions over these conditions evaluated both the models’ ability to interpolate (K_1_ and K_2_) and extrapolate (K_3_) responses to exoskeleton torques included in the training set. For each held-out condition (K_1_, K_2_, or K_3_), we trained the models using kinematic, EMG, and exoskeleton torque inputs from the zero-torque (K_0_) condition and the two non-zero-torque exoskeleton conditions not held out for validation. We evaluated each model by predicting output variables from the held-out exoskeleton condition using input data at an initial gait phase in the same condition. We compared prediction accuracies across held-out conditions.

To compare differences in performance across the three models, we identified differences in the models’ prediction accuracies using repeated-measures analysis of variance tests at a significance level of α = 0.05. When significant differences between models emerged, we identified pair-wise differences between models using post-hoc paired t-tests (α = 0.05) and a Holm-Sidak step-down correction for multiple comparisons [9, 42]. We report percent reductions in RRV values compared to the PV model and percent differences between the LPV and NPV models.

## V. RESULTS

The ankle exoskeletons had the largest impact on ankle kinematics, smaller impacts on knee and hip kinematics, and variable impacts on muscle activity (Fig. 2). Compared to the K_0_ condition, the peak ankle dorsiflexion angle during single-limb support decreased significantly in the K_2_ (36.7%) and K_3_ (40.0%) conditions (p < 0.020). Average integrated EMG increased slightly, but not significantly in the hamstrings and tibialis anterior (p > 0.066) in the K_2_ and K_3_ conditions compared to the K_0_ condition.

**Fig. 2.**
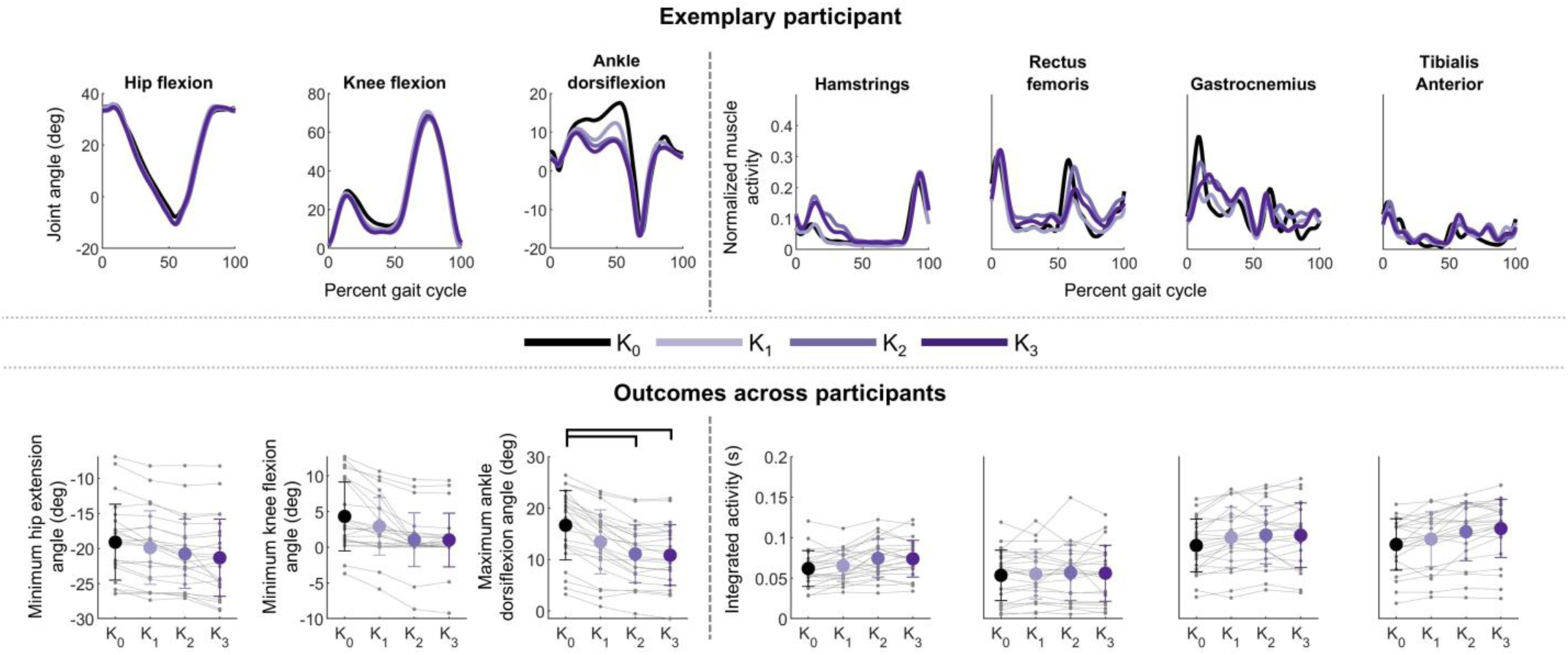
<u>Top</u>: Average kinematic (left) and EMG (right) data for one participant who exhibited large, repeatable responses to exoskeleton torque and high model prediction accuracies (P03). Black lines show the zero-torque condition (K_0_) that was subtracted from all conditions to reflect responses to exoskeleton torque. <u>Bottom</u>: Average (±1SD) kinematic and myoelectric responses for all participants in each torque condition. Brackets denote significant differences between exoskeleton conditions according to post-hoc paired t-tests (α = 0.05) and a Holm-Sidak step-down correction. Thin gray lines represent individual legs.

When validating on the held-out K_2_ condition, all three models predicted kinematic but not myoelectric responses to exoskeleton torque (Fig. 3, dashed lines). At a prediction horizon of Δ = 12.5% of a stride, the LPV model’s prediction accuracy at the ankle – where the largest responses to torque were observed – was 41.6 ± 16.0% more accurate than the PV model (p < 0.001) but not the NPV model (p = 0.130; Fig. 4; Table II). Similarly, the LPV model’s prediction accuracy at the hip was 41.7 ± 12.7% better than the PV model (p < 0.001). However, as prediction horizon increased, the average LPV and NPV model prediction accuracies of all outputs except the ankle approached those of the PV model. Changes in knee and hip kinematics were predicted more accurately than the baseline PV model for prediction horizons shorter than Δ = 18.75% of a stride (p < 0.001) in the LPV model and Δ = 12.5% of a stride (p < 0.001) in the NPV model (Fig. 5). At the ankle, the LPV model predicted kinematics 29.1–60.0% more accurately than the PV model for all prediction horizons (p < 0.001). The NPV model’s predictions were significantly more accurate than those of the PV model for all prediction horizons except 25.0% and 75.5–81.3% of a stride (p < 0.001).

**Fig. 3.**
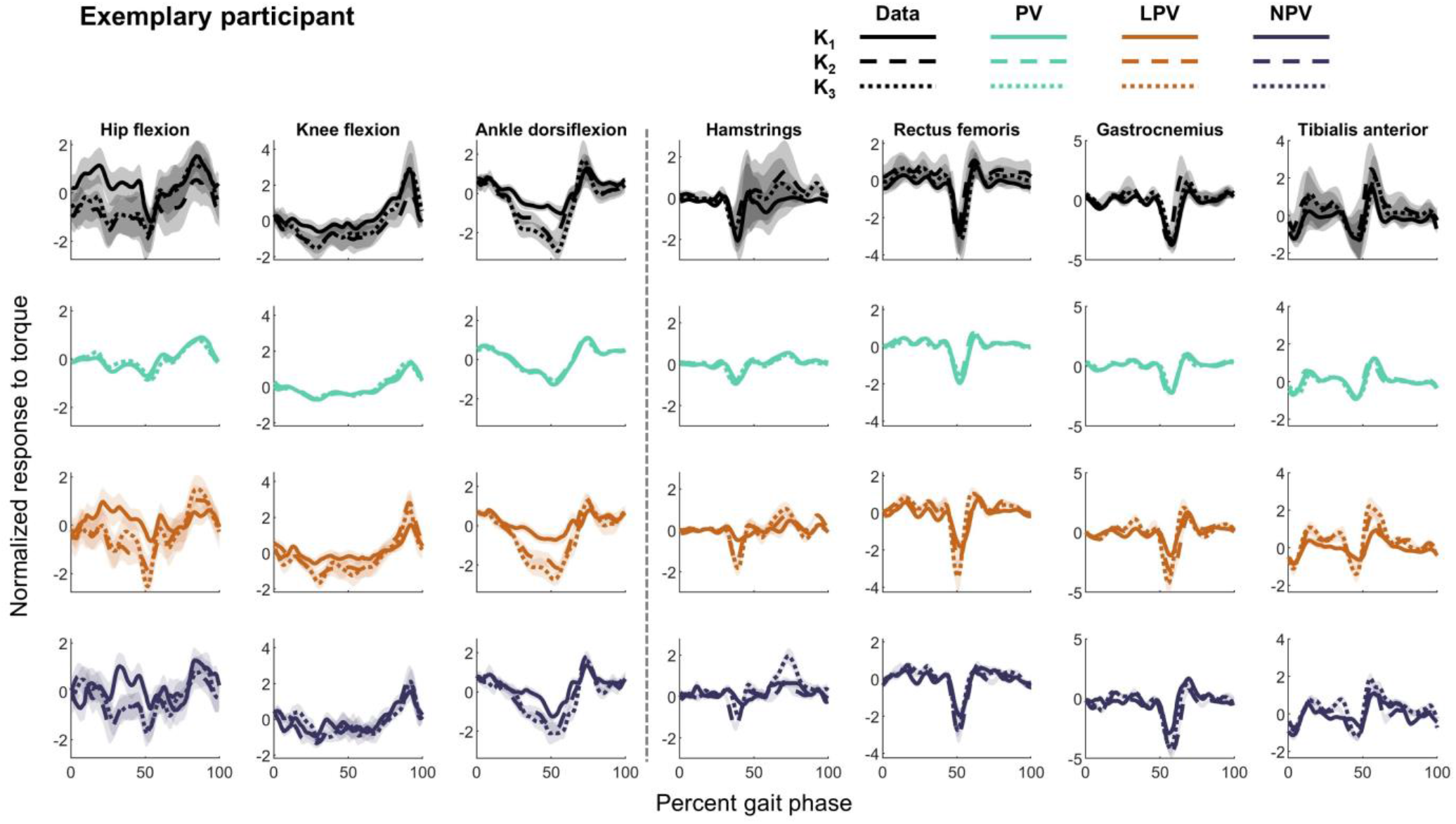
Kinematic and myoelectric experimental (black) and predicted (colors) responses to torque for one participant who exhibited large responses to the exoskeletons (P03). Predictions are shown for a prediction horizon of 12.5% of a stride for the PV (green), LPV (orange), and NPV (purple) models. The three held-out conditions are denoted with solid (K_1_), dashed (K_2_), and dotted (K_3_) lines. Lines represent the average (±1SD; shaded region) data and predictions over all gait cycles in the corresponding validation dataset. The experimental data show that the K_2_ and K_3_ responses to torque were more similar to each other than to the K_1_ response. The PV model predictions were similar across held-out conditions, while the LPV and NPV models scaled with exoskeleton torque. Full joint trajectories may be reproduced by rescaling and adding the average unperturbed gait cycle to the predictions. All comparisons used paired t-tests (α = 0.05) with a Holm-Sidak step-down correction for multiple comparisons.

**Fig. 4.**
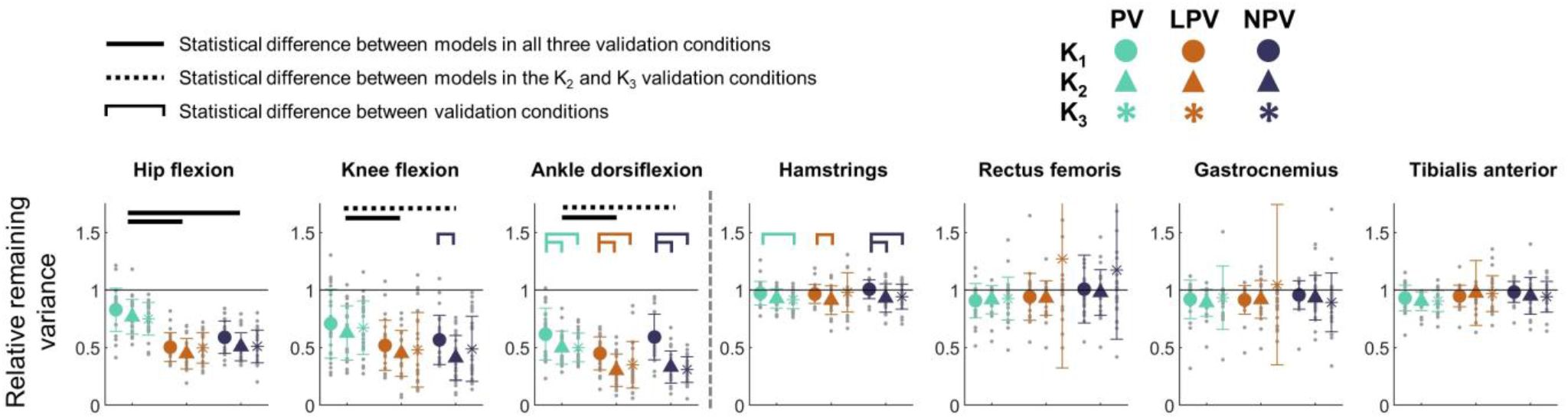
Average (±1SD) prediction accuracies for all participants and held-out conditions at a prediction horizon of 12.5% of a stride. Gray dots represent individual legs. Colored brackets denote statistically significant differences between held-out conditions for each model. Black horizontal bars denote significant differences between models across all three (solid) or two (dashed) held-out conditions. The large variance in the LPV model’s predictions of rectus femoris and gastrocnemius responses in the held-out K_3_ condition were due to bad predictions (RRV > 2) in a small number of legs. All comparisons used paired t-tests (α = 0.05) with a Holm-Sidak step-down correction for multiple comparisons.

**Table II:** Average (± 1SD) RRV values for kinematic and myoelectric predictions at a 12.5% prediction horizon.

| Output | PV | LPV | NPV |
| --- | --- | --- | --- |
| Ankle angle<sup>†‡</sup> | $0.50 \pm 0.14$ | $0.30 \pm 0.14$ | $0.33 \pm 0.14$ |
| Knee angle<sup>†‡</sup> | $0.62 \pm 0.24$ | $0.45 \pm 0.20$ | $0.41 \pm 0.19$ |
| Hip angle<sup>†‡</sup> | $0.77 \pm 0.15$ | $0.44 \pm 0.13$ | $0.51 \pm 0.12$ |
| Tibialis anterior | $0.90 \pm 0.08$ | $0.97 \pm 0.28$ | $0.95 \pm 0.15$ |
| Soleus | $0.86 \pm 0.21$ | $1.02 \pm 0.66$ | $1.04 \pm 0.64$ |
| Gastrocnemius | $0.89 \pm 0.12$ | $0.91 \pm 0.16$ | $0.93 \pm 0.19$ |
| Vastus medialis | $0.92 \pm 0.12$ | $0.93 \pm 0.15$ | $0.98 \pm 0.20$ |
| Rectus femoris | $0.87 \pm 0.21$ | $0.91 \pm 0.38$ | $0.94 \pm 0.42$ |
| Lateral hamstrings | $0.93 \pm 0.08$ | $0.91 \pm 0.13$ | $0.93 \pm 0.12$ |
| Gluteus medius | $0.95 \pm 0.08$ | $0.96 \pm 0.10$ | $0.98 \pm 0.10$ |

LPV = Linear phase-varying model; NPV = Nonlinear phase-varying model; PV = Phase-varying model
† Significant difference in prediction accuracy between the PV and LPV models
‡ Significant difference in prediction accuracy between the PV and NPV models

**Fig. 5.**
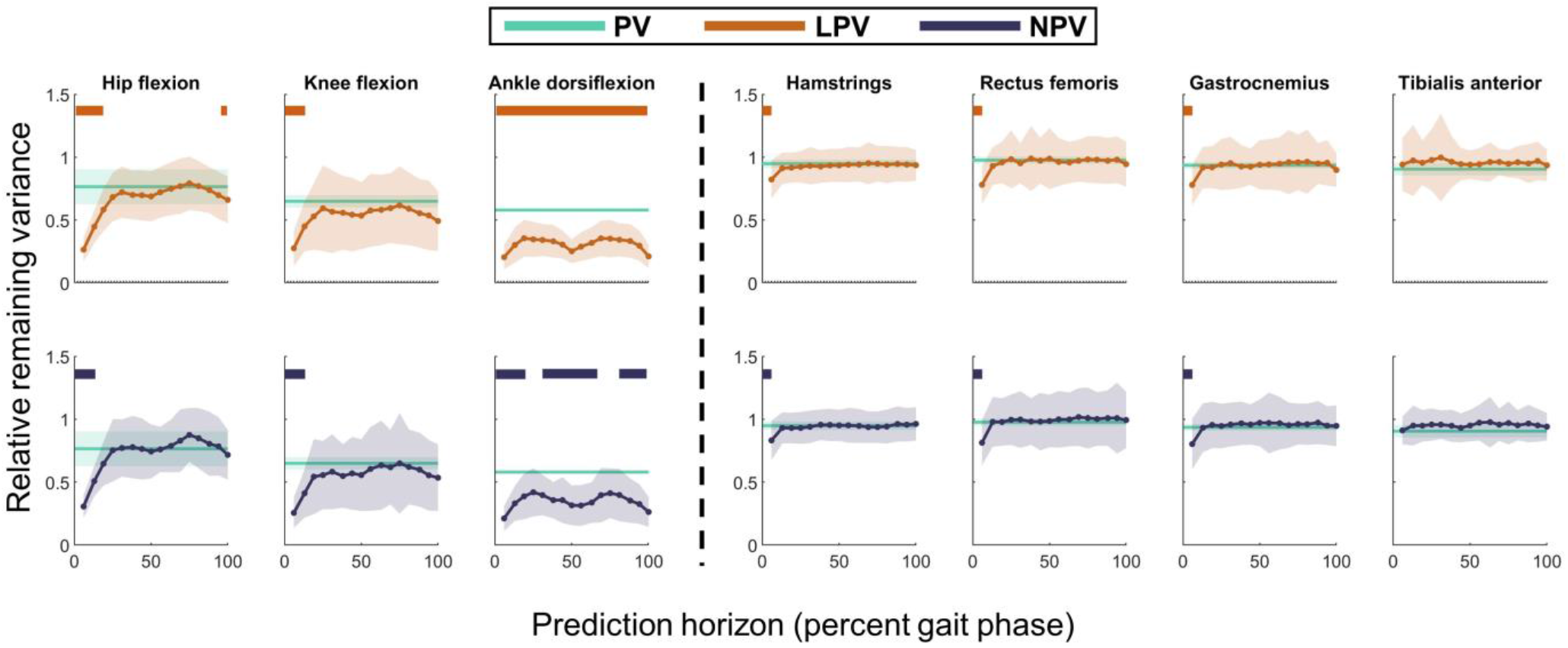
Prediction accuracy decreased with increasing prediction horizon. The PV model’s predictions (green) were constant across prediction horizons. Horizontal bars denote predictions that were significantly more accurate than the PV model. The LPV (top; orange) and NPV (bottom; purple) models’ prediction accuracies approached nearly constant values for prediction horizons beyond 25.0% of a stride for kinematic responses and 6.25% of a stride for myoelectric responses. The LPV and NPV models’ predictions of ankle kinematics remained more accurate than PV model predictions across almost all prediction horizons, while knee and hip kinematic predictions were similar to those of the PV model beyond 25.0% of a stride.

Predictions of myoelectric responses were poor (RRV ≈ 1.00) across all muscles and models, except at the shortest prediction horizon (Δ = 6.25%). At the shortest prediction horizon, both the LPV and NPV models’ predictions for the hamstrings, rectus femoris, and gastrocnemius were 10.7–15.0% more accurate than those of the PV model (p < 0.001; Fig. 5).

The LPV and NPV models’ prediction accuracies improved with increasing training set size. As expected, the PV model’s prediction accuracy was nearly constant across training set sizes (p > 0.005; Fig. 6). For a prediction horizon of Δ = 12.5% of a stride, the LPV model’s hip (RRV = 0.81) and knee (RRV = 0.78) prediction accuracies were significantly worse than RRV_full_ when using less than 50 strides of training data per exoskeleton condition (p < 0.001). Similarly, the NPV model’s hip and knee prediction accuracies approached RRV_full_ with approximately 50 strides of training data per condition (p < 0.001).The LPV model required more data – up to 150 strides per condition – for prediction accuracies to approach RRV_full_ at the ankle, gastrocnemius, and tibialis anterior (p < 0.001), though predictions were only 0.02–0.05 RRV points greater than RRV_full_ with 75 strides of training data per condition. The NPV model’s myoelectric prediction accuracies approached RRV_full_, in 25–75 strides of training data per condition (p < 0.001; Fig. 6).

**Fig. 6.**
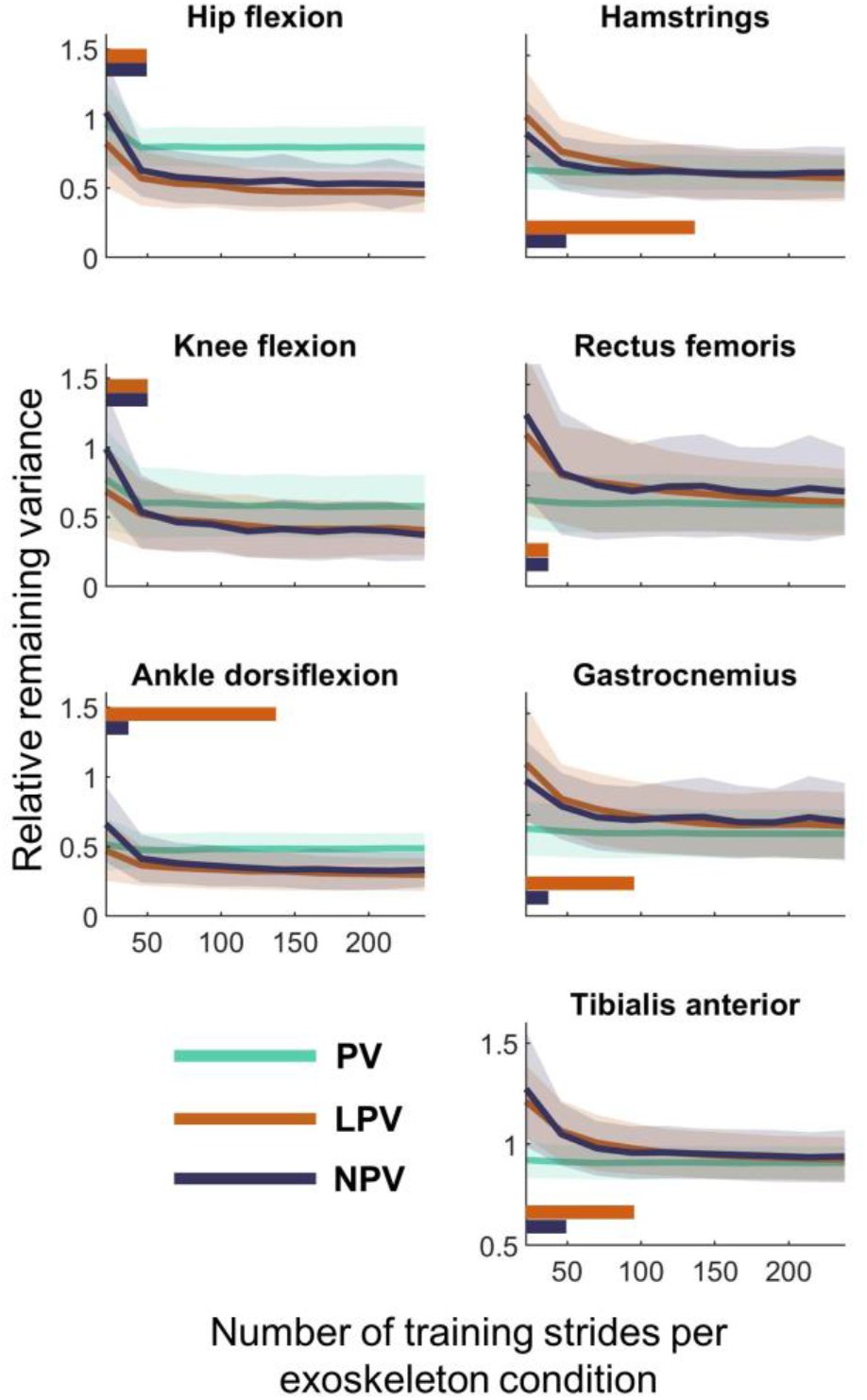
Average (±1SD; shaded region) prediction accuracy of kinematic (left) and myoelectric (right) outputs for the PV (green), LPV (orange), and NPV (purple) models over training set sizes ranging from 24 to 240 cycles (RRV_full_). Prediction accuracies were reported at a 12.5% stride prediction horizon. Orange (LPV) and purple (NPV) horizontal bars denote the training set sizes that yielded significantly worse predictions than those of the full training set. The PV model’s prediction accuracies were not significantly different from RRV_full_ at any training set size.

When validating on the held-out K_1_, K_2_, and K_3_ conditions, the LPV and NPV model predictions reflected experimental changes in response between conditions (Fig. 3). For all models at a 12.5% stride prediction horizon, predictions of responses in the held-out K_1_ condition (interpolation) were 0.10–0.28 RRV points at the ankle and 0.04–0.09 points in the hamstrings less accurate than predictions of the K_2_ or K_3_ datasets (p < 0.001). Conversely, no statistical differences in prediction accuracies of the held-out K_2_ (interpolation) and K_3_ (extrapolation) conditions were identified (Fig. 4). Improvements in kinematic prediction accuracy of the LPV model compared to the PV model were identified across the held-out K_1_, K_2_, and K_3_ conditions (p < 0.001). Differences between the NPV and PV models’ kinematic prediction accuracies in the held-out K_1_ condition did not reach significance at the knee or ankle.

## VI. Discussion

We evaluated the ability of subject-specific phase-varying models to predict kinematic and myoelectric responses to ankle exoskeleton torques during treadmill walking. When predicting across three exoskeleton torque conditions, both linear and nonlinear models predicted kinematic responses to exoskeletons without knowledge of the specific user’s physiological characteristics, supporting their potential utility as predictive tools for exoskeleton design and control. To our knowledge, this is the first study to predict kinematic and myoelectric responses to ankle exoskeletons using phase-varying models. Consistent with Floquet Theory and prior models of human locomotion, LPV models appear appropriate for predicting responses to exoskeleton torque over short prediction horizons, evidenced by its similar prediction accuracy to the more complex NPV model and improved prediction accuracy over the less complex PV model [25–27, 29].

The small and variable responses to exoskeleton torque exhibited by the unimpaired adults in this work highlight the challenge of altering kinematics with passive ankle exoskeletons. We found that even stiff exoskeletons (K_3_ = 5.08 Nm/deg) only altered ankle kinematics on average by six degrees and integrated muscle activity by 14%. These small changes may correspond to larger changes in joint powers or metabolic demands and indicate that the present study is a rigorous test case [1, 2, 5, 24]. Despite small changes in gait, the LPV model’s predictions explained more of the variance in kinematic responses to exoskeletons than the PV model, regardless of whether predictions interpolated (K_1_ and K_2_) or extrapolated (K_3_) relative to the training set. The LPV model’s ability to predict kinematics within and slightly beyond the available training data supports its potential utility for predicting responses to untested exoskeleton designs or control laws. However, predictions of the held-out K_1_ condition highlight the importance of selecting experimental conditions that encode complex responses to torque.

Our hypothesis that the LPV model would predict kinematic and myoelectric responses more accurately than the PV model and as accurately as the NPV model was partially supported. The LPV model’s kinematic and myoelectric predictions were more accurate than those of the PV model only for prediction horizons less than 18.75% and 6.25% of a stride, respectively, but the LPV and NPV models exhibited similar prediction accuracies across prediction horizons. The LPV and NPV models’ similar predictions support research demonstrating that nonlinear spring-loaded inverted pendula (SLIPs) have similar predictive accuracy to linear models of human movement [25]. Compared to a nonlinear SLIP, the NPV model’s feedforward neural network imposed fewer restrictions on model structure and enabling greater differences in prediction accuracy compared to a linear model. Therefore, the similarity of LPV and NPV model predictions supports the extension of Floquet Theory to gait with exoskeletons and indicates that, for rhythmic locomotion at a constant speed over level ground, linear phase-varying models have sufficiently complex structure to predict kinematic responses to exoskeletons [25–27, 29].

We observed comparable kinematic prediction accuracy to studies using physics-based and data-driven models of locomotion. Maus et al. evaluated multiple models’ abilities to predict center-of-mass height during running and reported accuracies ranging from RRV ≈ 0.15 at a 15% prediction horizon to RRV ≈ 0.85 beyond an 80% stride prediction horizon, for an exemplary participant [25]. Within a similar range of prediction horizons, the LPV model predicted kinematics across participants with average accuracies ranging from (0.30 < RRV < 0.45) at a 12.5% prediction horizon and (0.34 < RRV < 0.77) at an 81.3% prediction horizon. Similarly, Drnach et al. [43] used a hybrid linear model to predict response to functional electrical stimulation, reporting median RRV values (transformed from a fitness score) ranging from approximately 0.11-1.04. However, the average unperturbed gait cycle was not subtracted from the data before computing the fitness score in [43]. The average unperturbed cycle accounts for a substantial portion of the variance in the perturbed signals, providing a less conservative prediction accuracy statistic than the RRV presented here. For example, if the unperturbed cycle had not been subtracted from the data in the present study, the LPV model’s ankle predictions for one participant who exhibited large responses to torque would be RRV = 0.08 rather than the more conservative 0.21 reported. Comparable prediction accuracies to prior work indicate that phase-varying models are potentially useful predictive tools for locomotion with ankle exoskeletons and may have similar predictive power to physics-based models of locomotion.

The convergence of LPV and NPV models’ prediction accuracies to an approximately constant value at large prediction horizons (e.g. RRV_LPV_ ≈ 0.70 for knee kinematics at Δ > 25.0% of a stride) may be useful when selecting measurements for device design or control. The LPV and NPV models’ kinematic prediction accuracies decreased rapidly from 6.25% to 18.75% stride prediction horizons, before reaching an approximately constant value. Ankle predictions remained better than those of the PV model across prediction horizons. Higher prediction accuracy at the ankle was unsurprising due to large responses to exoskeletons and the ankle’s direct piecewise-linear relationship to passive exoskeleton torque. Since we trained on multiple exoskeleton conditions, the dynamics predicting future ankle kinematics are higher-dimensional than the simple exoskeleton torque-ankle angle relationship, suggesting that accurate predictions of ankle kinematics over large prediction horizons are likely for powered exoskeletons as well. Unlike the ankle, hip and knee kinematics were indirectly impacted by exoskeleton torque and their RRV values approached those of the PV model for prediction horizons above 18.75% of a stride. This result indicates that stride-specific initial posture and exoskeleton torque were predictive of indirect exoskeleton impacts on kinematics only for short prediction horizons. At large prediction horizons, measurements at an initial phase did not, on average, improve predictions of future posture. However, some participants’ hip and knee kinematics were predicted up to 0.30 RRV points more accurately by the LPV and NPV models than the PV model across prediction horizons, suggesting that the prediction horizon at which stride-specific measurements no longer improve predicted responses to exoskeletons depends on the magnitude of the individual’s response. The LPV and NPV models’ accurate predictions over short prediction horizons make them primarily useful for exoskeleton control [10, 11]. For individuals that exhibit large responses to exoskeletons, however, LPV model-based predictions over stance may inform passive exoskeleton parameter selection. Guided adaptation and extended practice sessions [1, 17] or powered ankle exoskeletons [5, 6] may elicit larger responses than those observed in this study and increase the maximum prediction horizons at which measurements at an initial posture improve predicted responses to torque, potentially expanding the settings in which model predictions are useful.

A major limitation of all three models was their inability to predict myoelectric responses. The LPV and NPV models predicted myoelectric signals more accurately than the PV model only for the shortest prediction horizon (Δ = 6.25%). While exoskeleton torque and stiffness are known to impact average plantarflexor activity, we found that the average unperturbed gait cycle accounted for only 30-60% of the variance in the K_2_ data, compared to 60-95% in kinematic signals [1, 9, 12]. Consequently, poor prediction accuracy may be partially attributed to small changes in muscle activity between the exoskeleton conditions. Alternatively, kinematic and myoelectric input variables may fail to encode nonlinear musculotendon dynamics, which are impacted by ankle exoskeletons, between the initial and final phases [21, 40]. Studies predicting muscle activity using physiologically-detailed models accounted for 60-99% of the variance in myoelectric signals, though they evaluated predictions on unperturbed walking conditions only [44, 45]. Still, the difference in prediction accuracy between the phase-varying models and physiologically-detailed models indicates that encoding musculotendon dynamics in the input variables is likely needed to improve myoelectric predictions for data-driven phase-varying models and represents an interesting area of future research.

Another limitation of subject-specific data-driven models, compared to physiologically-detailed models, is the amount of training data required to predict changes in gait with exoskeletons, which impacts models’ utility in settings where minimizing data collection duration is critical to mitigating physical and logistical burdens on participants and families, such as in clinical gait laboratories. Improvements in prediction accuracy of the LPV and NPV models were small beyond 75–100 strides of training data per exoskeleton condition. The LPV model required more training data at the ankle, but a similar amount at the hip and knee to that used by Drnach et al., who trained a hybrid linear model using 45 seconds of data across two experimental conditions [43]. For unimpaired, steady-state locomotion, data-driven linear models appear to require 75–125 strides of training data per condition, which supports their feasibility only in gait analysis settings with treadmills or long walkways [6, 7]. Additional dimensionality reduction, such as via sparse regression, may reduce the LPV model’s complexity and demand for training data [25, 31, 46]. However, when only one training condition or a few strides are collected, as is standard in clinical gait analysis, phase-varying model predictions will be poor and physiologically-detailed or population-specific models may generate more accurate predictions [8, 19, 44, 45].

Subject-specific data-driven phase-varying models of gait with exoskeletons have benefits and limitations compared to predictive musculoskeletal models. While we investigated only a specific subset of phase-varying models, we showed that this class of model can predict kinematic responses to exoskeletons without detailed knowledge of the physiological and neuromuscular factors influencing responses to exoskeletons. Conversely, uncertainty in the mechanisms driving complex responses to exoskeletons may limit physiologically-detailed models’ accuracy [13, 24]. While predictive musculoskeletal models may generate “what-if” predictions without experimental data, data may be needed to specify initial postures and tune subject-specific parameters. Phase-varying models can similarly perform subject-specific “what-if” predictions when application-specific training data are available. Unlike physiologically-detail models, this and prior work exemplify phase-varying models’ ability to take arbitrary measurements as inputs, enabling their application using a range of experimental resources [25, 26, 31]. Extending data-driven predictions to “what-if” scenarios and improving predicted myoelectric responses to exoskeletons, combined with analytical tools for phase-varying systems(e.g. [31]), may facilitate prediction and analysis of individualized exoskeleton impacts on gait mechanics and motor control.

## VII. Conclusion

To our knowledge, this is the first study to predict subject-specific responses to ankle exoskeletons using phase-varying models. Without making assumptions about individual physiology or motor control, an LPV model predicted short-time kinematic responses to bilateral passive ankle exoskeletons, though predicting myoelectric responses remains challenging. Results support the utility of LPV models for studying and predicting response to exoskeleton torque. Improving data-driven models and experimental protocols to study and predict myoelectric responses to exoskeletons represents an important direction for future research. Modeling responses to exoskeletons or other assistive devices using a phase-varying perspective has the potential to inform exoskeleton design for a range of user groups.

## VIII. Author Contributions

MCR was involved in the conception and design of the study, carried out exoskeleton design and fabrication, data collection, data preprocessing, statistical analyses and data interpretation, and drafted the manuscript. BSB was involved in the conception and design of the study and carried out model development and coding, contributed to the initial manuscript draft, and critically revised the manuscript. SAB critically revised the manuscript and acquired funding for this work and was involved in the conception and design of the study, model development, and data interpretation. KMS critically revised the manuscript, acquired funding for this work, and was involved in the conception and design of the study and data interpretation.

## IX. Data Accessibility

All experimental datasets and code using in this study are freely available and can be accessed on https://simtk.org/projects/ankleexopred.

## X. Ethics

This study was approved by the University of Washington Institutional Review Board #47744. All research participants provided informed consent prior to participating in the study, obtained by MCR.

