## Supplement - S1 for "Predicting walking response to ankle exoskeletons using data-driven models"

### SUMMARY OF CONTENTS

This work evaluated the ability of phase-varying models to predict kinematic and myoelectric responses to bilateral ankle exoskeleton torque during steady-state locomotion in unimpaired adults. This document provides additional theory and results supporting modeling decisions. The following sections discuss:

I. Exoskeleton stiffness and torque profiles as inputs to the phase-varying models
II. Ground reaction force data and predicting joint dynamics
III. Individual responses to varying prediction horizon

**Data availability**: Datasets and code are freely available at *https://simtk.org/projects/ankleexopred*.

### I. EXOSKELETON STIFFNESS AND TORQUE PROFILES AS INPUTS TO PHASE-VARYING MODELS

We used passive exoskeleton torque profiles, rather than stiffness as inputs to our linear and nonlinear phase-varying models. We discretized torque profiles between the initial and final phases (10 samples per leg) as inputs to the model. However, based on our fitting procedure – using multiple exoskeleton conditions to fit a single model – the models do not simply predict ankle angle purely as a function of torque.

As noted in the main manuscript, using stiffness rather than torque as a model input is possible, but lacks physical or theoretical basis, whereas torque represents the mechanical interaction between the exoskeleton and the user. Our decision to use exoskeleton torque samples was driven by the theoretical support that the future state of a discrete-time linear model, such as the linear phase-varying (LPV) model, is a linear function of the initial state and the entire input history between initial and final times [1]. Similar theory supports stiffness being insufficient as an input variable to the LPV model: While torque is a linear function of stiffness and ankle kinematics, ankle kinematics are a nonlinear function of phase, indicating that using stiffness as a model input is not equivalent to using torque as an input. However, since the nonlinear phase-varying (NPV) model may use nonlinear functions of stiffness, it is possible that using stiffness as an input would generate similarly accurate predictions to the torque-based NPV model.

We compared the prediction accuracy of stiffness-based LPV and NPV models to torque-based models using the $K_2$ validation condition. The torque inputs used ten time-history samples per leg, while the stiffness condition used a single constant stiffness value per exoskeleton condition as an input. We expected prediction accuracies to be lower in the stiffness-based models, due to theoretical support against stiffness as an input and the added information in the torque samples. Indeed, The LPV model’s predictions using the stiffness input were 0.16 ± 0.13 RRV points worse than with torque inputs at the ankle ($p < 0.001$) according to paired t-tests with Holm-Sidak correction for multiple comparisons at a significance level of α = 0.05 (Fig. S1) [2]. The NPV model was more sensitive to input type than the LPV model and its predictions were worse at the hip (0.19 ± 0.19 RRV points), knee (0.37 ± 0.21 RRV points), and ankle (0.41 ± 0.16 RRV points) when using stiffness as an input (all $p < 0.001$), compared to using torque inputs. The reduced prediction accuracy in the stiffness-based LPV and NPVs model does not support the use of stiffness as an input variable to phase-varying models.

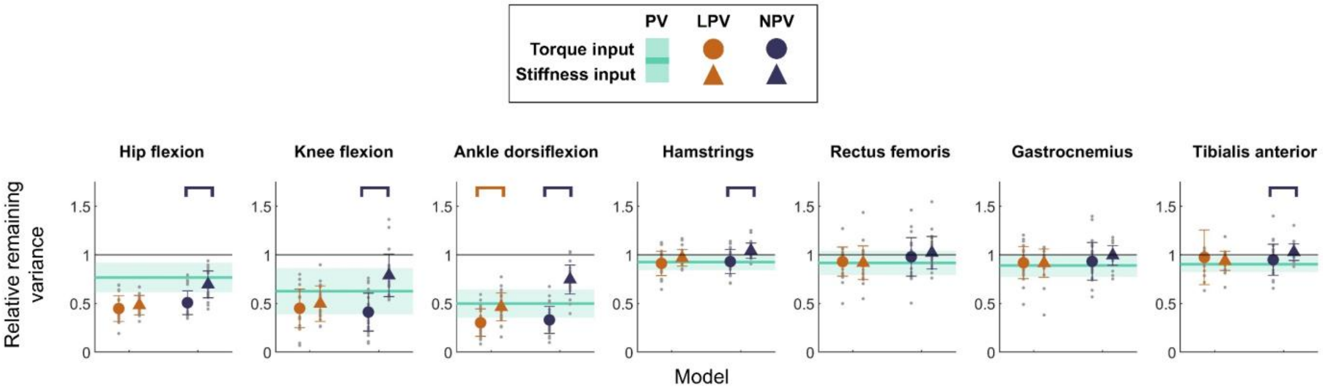


**Fig. S1. A comparison of prediction accuracies for stiffness (triangles) and torque (circles) inputs in the $K_2$ validation condition. Averages (±1SD) are shown for the LPV and NPV models. The PV model shows the average (±1SD) as a solid line (shaded region). The PV (green) model was agnostic to inputs and was constant across input conditions. Brackets denote significant differences between torque and stiffness prediction accuracies according to paired t-tests with Holm-Sidak correction for multiple comparisons at a significance level of α = 0.05.**

### II. GROUND REACTION FORCE DATA AND PREDICTING JOINT DYNAMICS

Data-driven phase-varying models can accept arbitrary input and output variables, which makes predicting GRFs or joint moments an interesting area of future research. Our dataset was inappropriate for predicting dynamic variables for two reasons:

1. Some participants frequently stepped on both belts of the split-belt treadmill with one foot, causing incorrect estimates of the center of pressure and GRFs corresponding to each leg. Sporadic inaccuracies in the GRF and center of pressure data may result in model fits and predictions being highly sensitive to which gait cycles are included in the training and validation datasets.

2. Changes in GRFs between exoskeleton conditions were small (Fig. S2). Although we observed statistical differences ($p < 0.05$; paired t-tests) for the anterior-posterior and vertical GRFs, the differences between conditions were small: (Cohen's $d < 0.20$) for all GRF directions, and frequently less than 0.05 [3]. Moreover, the zero-stiffness baseline exoskeleton condition accounted for 99% [97.5%, 99.4%] (median [IQR]) of the variance across all GRF signals. Consequently, the LPV or NPV models' predictions would at most account for an additional three percent of the variance in the GRF data.

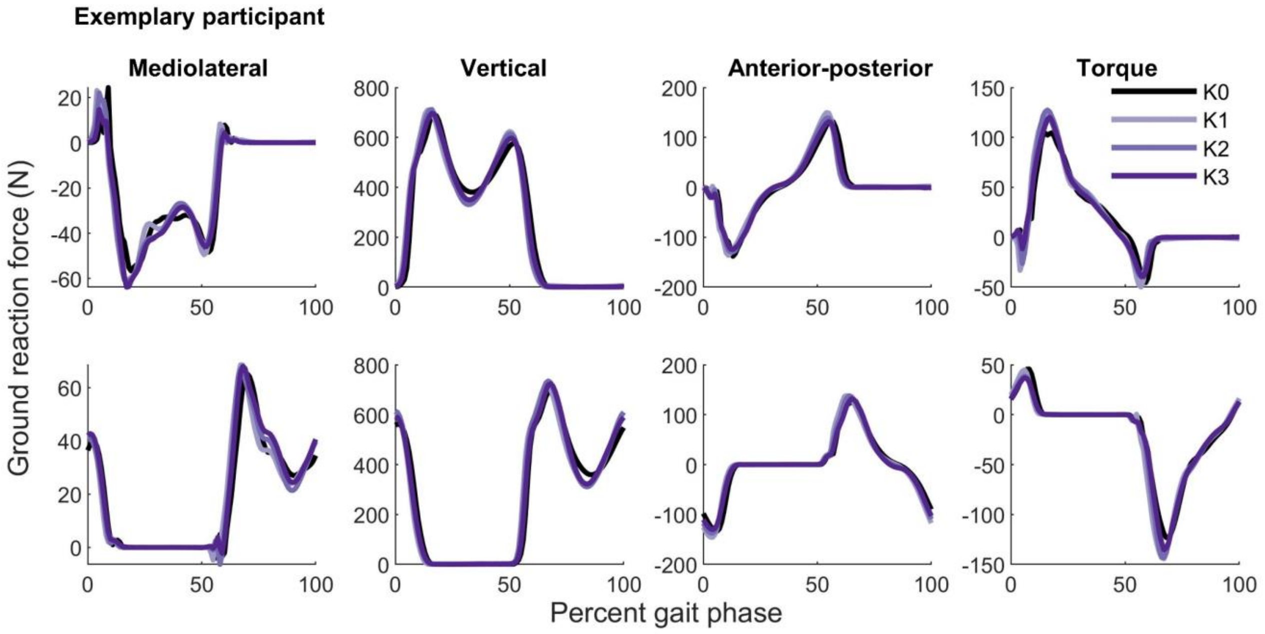


**Fig. S2. Exemplary average ground reaction force across the four exoskeleton conditions data for one participant (P03) that exhibited a large kinematic response to ankle exoskeletons. The signals represent the data averaged over all gait cycles of each condition. Other participants' ground reaction force data exhibited similar or smaller changes between exoskeleton conditions.**

### III. INDIVIDUAL RESPONSES TO VARYING PREDICTION HORIZON

While average model predictions approached those of the PV model at large prediction horizons, the LPV and NPV models explained some of the variance in participants' gait kinematics (Fig. S3) and muscle activity (Fig. S4) for prediction horizons spanning the entire gait cycle.

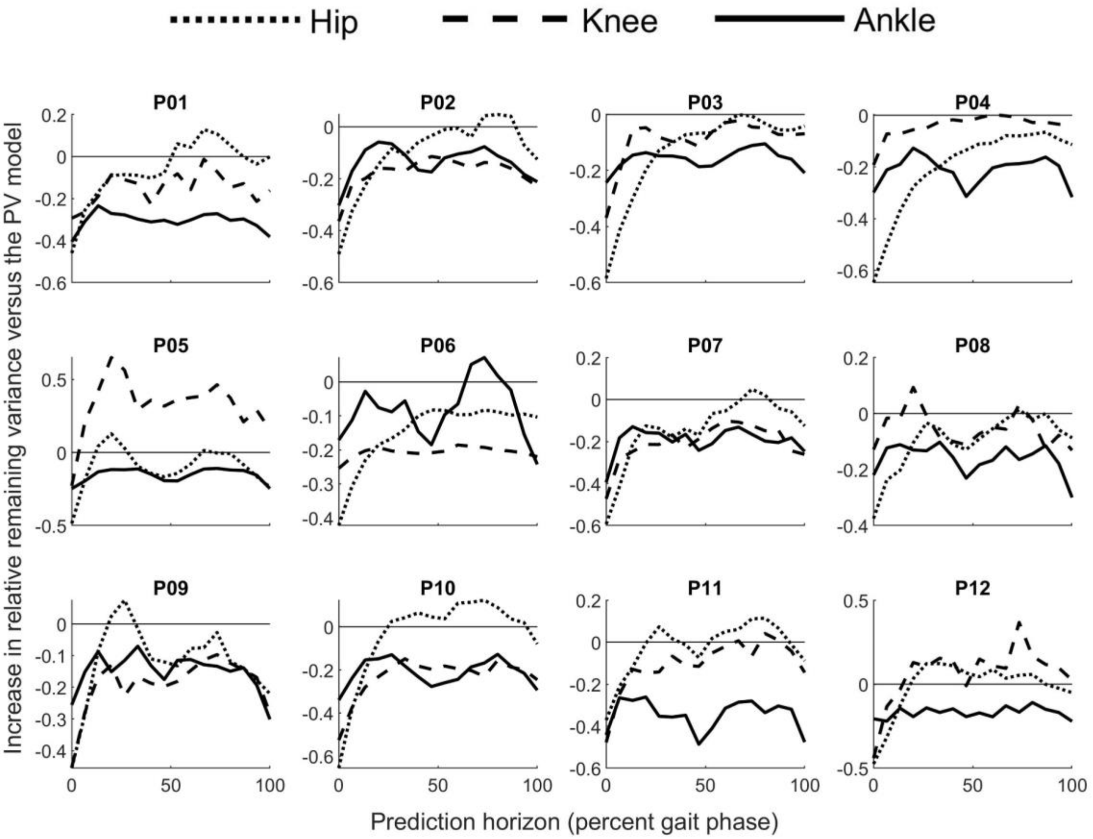


**Fig. S3. Change in relative remaining variance (RRV) of the LPV model compared to the PV model across prediction horizons for each participant's kinematic responses to torque in the $K_2$ validation condition. The RRV values were averaged across legs. An RRV value < 0 implies that the prediction is better than guessing the average response at each gait phase (*i.e.* the PV model).**

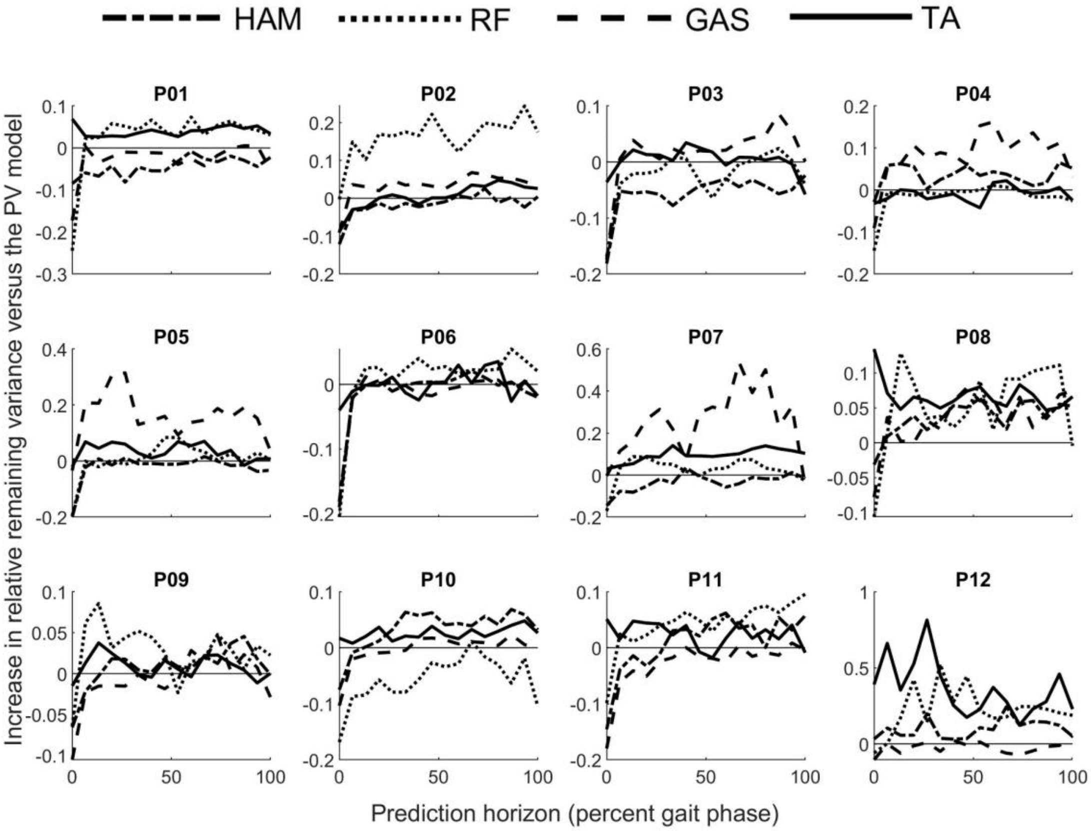


**Fig. S4. Change in relative remaining variance (RRV) of the LPV model compared to the PV model across prediction horizons for each participant's myoelectric responses to torque in the $K_2$ validation condition. The RRV values were averaged across legs. An RRV value < 0 implies that the prediction is better than guessing the average response at each gait phase (*i.e.* the PV model).**
