## Supplement - S2 for "Predicting walking response to ankle exoskeletons using data-driven models"

SUMMARY OF CONTENTS

This work evaluated the ability of phase-varying models to predict kinematic and myoelectric responses to bilateral ankle exoskeleton torque during steady-state locomotion in unimpaired adults. This document provides additional details of the experimental setup and additional information to facilitate replication of our work. The following sections discuss:

I. Experimental setup & participant descriptions
II. Computing joint kinematics, and estimating exoskeleton torque
III. Model algorithms and explanations of the processing pipeline
IV. Additional information on prediction accuracy evaluation

**Data availability**: Datasets and code are freely available at *https://simtk.org/projects/ankleexopred*.

### I. EXPERIMENTAL SETUP AND PARTICIPANTS

All participants were unimpaired adults (12 unimpaired adults (6 female / 6 male; age=23.9±1.8 yrs; height=1.69±0.10 m; mass=66.5±11.7 kg; Table SI).

TABLE SI: PARTICIPANT CHARACTERISTICS

| Subjects | Gender (M/F) | Age (years) | Mass (kg) | Height (m) | Walking speed (m/s) |
|---|---|---|---|---|---|
| P01 | M | 25 | 73.5 | 1.78 | 1.36 |
| P02 | F | 24 | 61.2 | 1.65 | 1.40 |
| P03 | F | 20 | 49.9 | 1.60 | 1.30 |
| P04 | F | 22 | 65.8 | 1.73 | 1.50 |
| P05 | F | 25 | 55.0 | 1.55 | 1.30 |
| P06 | F | 23 | 49.4 | 1.52 | 1.30 |
| P07 | M | 27 | 59.9 | 1.68 | 1.35 |
| P08 | M | 23 | 77.1 | 1.73 | 1.55 |
| P09 | M | 24 | 80.7 | 1.84 | 1.40 |
| P10 | M | 25 | 73.9 | 1.73 | 1.20 |
| P11 | M | 25 | 83.9 | 1.83 | 1.40 |
| P12 | F | 24 | 68.0 | 1.70 | 1.22 |

M = Male;  F = Female ; kg = kilograms; m = meters; m/s = meters per second

#### *A. Data collection protocol*

Participants underwent two sessions: The first session – exoskeleton fitting and practice – ensured that the appropriately sized footwear was selected, that the exoskeleton cuff was at a comfortable height, and any points of discomfort were adequately padded. Participants were instructed to stop if they experienced pain or modified their gait pattern to avoid discomfort. Each participant performed 20 minutes of walking practice under three exoskeleton stiffness conditions (K0, K1, K3), spanning the range of conditions used during data collection. Participants walked at three nondimensional speeds (0.35, 0.45, 0.55) and three cadences (100, 120, and 140 steps/minute), set by a metronome, for one minute each. The ordering of the speed and cadence conditions was (1) middle, (2) high, (3) low. Nondimensional speed, $\hat{v}$, was defined as $\hat{v} = \frac{v}{\sqrt{gL}}$, where $v$ is the walking speed in meters/second, $g$ is the gravitational constant, and $L$ is the leg length measured from the lateral malleolus of the ankle to the anterior superior iliac spine point on the pelvis (**Fig. S1**) [1]. Walking speeds during practice ranged from 0.88 m/s to 1.67 m/s. Following the practice session, each participant selected a preferred walking speed corresponding to a "pace that they could comfortably sustain for 60 minutes while walking with the exoskeletons." Beginning at the intermediate nondimensional speed, the treadmill speed was changed according to the participant's requests (faster/slower), until the participant identified a preferred speed. If a participant did not explore speeds above/below their selected speed, a second iteration was performed, beginning at a speed below/above the selected speed to encourage exploration of walking speeds during the selection process, which is known to influence gait pattern selection [2]. All participants selected walking speeds within the range of nondimensional speeds used during practice ($0.40 \leq \hat{v}_{selected} \leq 0.53$).

During the second session, participants walked at their previously-selected speed for six minutes per exoskeleton condition. The last four minutes of each condition were recorded. Exoskeleton conditions were randomized. Participants rested for approximately five minutes between trials. Before the first walking condition, we performed a static calibration while the participant stood with each foot on an individual force plate to measure the participant's mass with the exoskeletons. Recalibrations were performed when participants used the restroom or drank water.

*B. Marker and electromyography sensor placement*

Data were collected during the second session. We used a modified Helen Hayes marker set for motion capture and placed electromyography (EMG) sensors (Delsys Inc, Natick USA) according to SENIAM guidelines [3, 4]. The EMG sensors were placed bilaterally on the gluteus medius, rectus femoris, vastus medialis, biceps femoris, medial gastrocnemius, soleus, and tibialis anterior. Marker trajectories were collected using a 10-camera infrared motion capture system (Qualisys AB, Göteborg, SE). Participants walked on an instrumented split-belt treadmill (Bertec Inc, Columbus OH) and were instructed to walk naturally. Marker trajectories were sampled at 120Hz and EMG data were sampled at 2000Hz. All data were downsampled to 120Hz before exporting for processing and analysis.

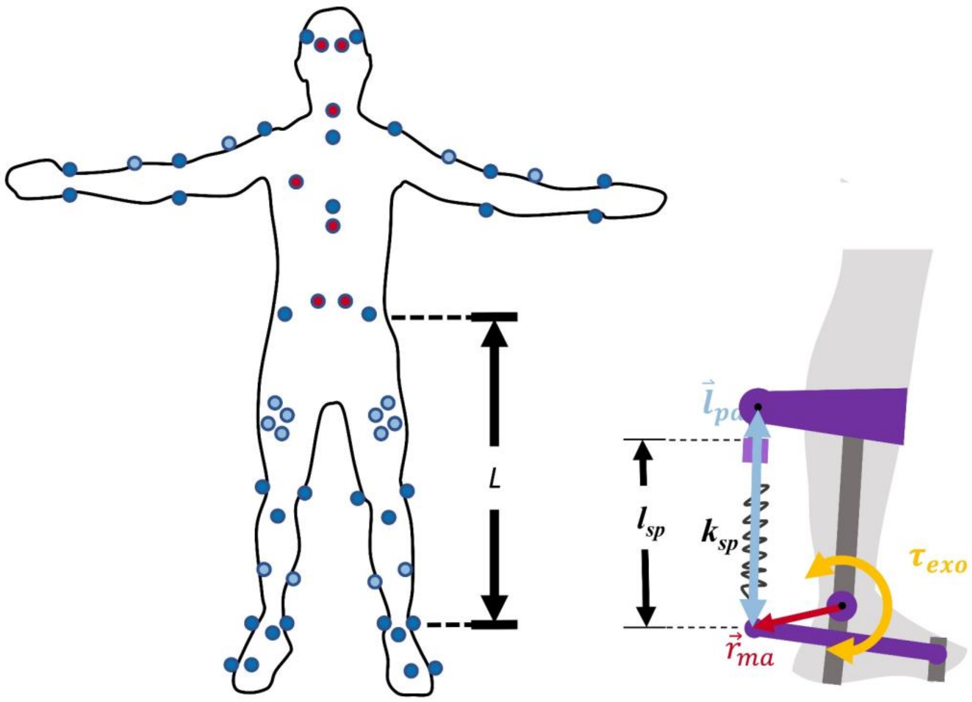


**Fig. S1.** **<u>Left</u>: A modified Helen Hayes marker set used in this study [3]. The depicted participant is facing forward. Dark blue markers were visible from an anterior view of the participant and were placed on bony landmarks on the body. Light blue markers were tracking markers and were also visible from an anterior view but were not placed on bony landmarks. Red markers were visible from a posterior view of the participant and were placed on bony landmarks. The distance *L* represents leg length. <u>Right</u>: A depiction of the exoskeleton components as listed in Eqn. 1. The exoskeleton torque, $\tau_{exo}$, is determined by $l_{pd}$, the distance between the proximal and distal moment arms, $l_{eq}$, the equilibrium length of the spring cable, the spring stiffness, $k_{sp}$, and the moment arm vector between the spring insertion and the ankle joint, $r_{ma}$.**

#### II. Data preprocessing

##### A. Computing joint kinematics

Marker trajectories were low-pass filtered at 6 Hz using a fourth-order zero-lag Butterworth filter. We converted marker trajectories to joint kinematics using OpenSim 3.3 [5]. First, we scaled a generic 29 degree-of-freedom skeletal model to each participant’s skeletal geometry [6]. The subtalar, metatarsophalangeal, wrist flexion, and wrist deviation degrees of freedom were locked. We ensured that root-mean-squared marker errors were less than 1 cm and maximum marker errors were less than 2 cm. Joint kinematics were then computed using OpenSim’s inverse kinematics algorithm. The inverse kinematics algorithm identifies joint angle trajectories to minimize error between model markers, which are fixed on rigid body segments of the model, and the experimental marker trajectories. The knee joint range of motion was increased to permit up to five degrees of hyperextension for participants who appeared to hyperextend their knee during walking. We evaluated model quality using experimental marker errors, in line with best-practices [7]. Root-mean-squared marker errors were less than 2 cm and maximum marker errors were less than 4 cm for each trial.

##### *B. Estimating exoskeleton torque profiles*

For a passive exoskeleton, the torque, $\tau_{exo}$, depended approximately linearly on the user's ankle kinematics, $\theta_{ankle}$, exoskeleton equilibrium angle, $\theta_{eq}$, and exoskeleton's rotational stiffness, $k_{exo}$ (Eqn 1) as:

$$\tau_{\mathrm{exo}}(\mathrm{t}) = \begin{cases} -\mathrm{k}_{\mathrm{exo}}\left(\theta_{ankle}(t) - \theta_{eq}\right), & \theta_{ankle} \geq \theta_{eq} \\ 0 & \theta_{ankle} < \theta_{eq} \end{cases} \quad (1)$$

Unlike many clinical exoskeletons, whose torque profiles are smooth functions of ankle angle, torque in the exoskeletons used in this work were piecewise-smooth functions of ankle angle, providing plantarflexion assistance similar to other experimental devices (Fig. S1) [8, 9].

The linear approximation in Eqn 1 was deemed sufficient based on calibration trials with pilot datasets, during which tension load cells (Omega Engineering, Norwalk, CT) were attached in series to each exoskeleton spring (Eqn 2). The measured torque from the load cell, $\tau_{exo}$, was,

$$\tau_{\mathrm{exo}}(\mathrm{t}) = \vec{r}_{\mathrm{ma}}(\mathrm{t}) \times \left[-k_{sp}\left(\vec{l}_{pd}(t) - l_{eq}\right)\right], \quad (2)$$

where $l_{pd}$ is the distance the proximal and distal spring attachment points, and $l_{eq}$ is the equilibrium length of the spring cable, which we estimated by having the subject dorsiflex until they felt the spring engage with the exoskeleton raised off the ground. The spring stiffness is denoted by $k_{sp}$. The cross product of the spring force vector and the vector between the moment arm at the distal end of the spring cable and the exoskeleton ankle joint, $r_{ma}$, defined the exoskeleton torque. Preliminary analyses showed small differences between torque profiles estimated from the nonlinear torque-ankle angle relationship (Eqn 2) and the linear relationship (Eqn 1) that we employed in our analysis.

#### III. Phase-varying model algorithms

We used hip flexion angles of the right and left limbs as phase variable inputs to the Phaser algorithm [10], which is similar to [11]. Phase estimates were used as the variable in the Fourier series representation of the phase-varying (PV) and linear phase-varying (LPV) models to cluster subsets of the time series data when fitting the LPV model and as inputs to the nonlinear phase-varying (NPV) model. We denoted phase as $\phi \in \mathbb{R}^{T}$, where $T$ denotes the number of samples in the training set. Each model had N = 20 outputs. The LPV and NPV models used the same M = 80 inputs, though the NPV model also took phase as an input.

##### *A. Phase-varying model*

The PV model, $F_{pv}^{w}$, is purely a function of phase and was represented in this work using a Fourier Series of order H=7 (Eqn. 3). The Fourier Series was parameterized by matrix coefficients $w_j, j \in \{1,2,\ldots,2H\}$, which were fit using a least-squares approximation of each trajectory.

$$F_{pv}^{w} = \frac{1}{2}w_0 + \sum_{h=1}^{H} w_{2h-1}\cos(h\phi) + \sum_{h=1}^{H} w_{2h}\sin(h\phi) \tag{3}$$

##### *B. Linear phase-varying model*

The LPV model is linear with respect to the inputs, $X(\phi)$, but nonlinear with respect to phase and has the form $X(\phi+\Delta)=A_{\phi,\Delta}X(\phi)$, where $\Delta$ denotes the lookahead window length. The LPV takes in a phase, $\phi$, and returns affine matrices $A_{\phi,\Delta}\in\mathbb{R}^{N\times M+1}$. For 64 phases over the cycle, we fit discrete maps $A_{\phi_l,\Delta}: X(\phi_l)\rightarrow X(\phi_l+\Delta)$ using a weighted least-squares regression with a Gaussian weighting scheme to penalize samples based on proximity to the phase $\phi_l$. To generate a continuous representation of the 64 discrete maps, $F_{LPV,\Delta}^{w}$, we parameterized each element of the maps using a seventh-order Fourier Series as a function of phase (Eqn. 3). We used the same least-squares approximation as with the PV model. This fitting procedure was repeated independently for each lookahead window.

##### *C. Nonlinear phase-varying model*

The NPV model used a fully-connected three-layer feedforward neural network to represent the response to torque. The model's hidden layer was 128 units wide – greater than the number of inputs to avoid enforcing a reduced-order representation of the input-output relationships – and used a softsign activation function [12, 13]. The NPV took the 80 inputs plus phase to predict the same 20 outputs as the PV and LPV models. We implemented the NPV model in the Keras Python framework, using RMSprop gradient descent. The NPV model's hyperparameters included interconnection weights and biases.

#### IV. Evaluating model prediction accuracy

##### *A. Relative Remaining Variance*

The Relative Remaining Variance (RRV) outcome used in this study is defined as

$$RRV=\frac{var\left(Y-\hat{Y}\right)}{var(Y)}, \tag{4}$$

$$0<RRV<\infty \tag{5}$$

where *var* denotes the variance in the each output, $Y$ denotes the test data, and $\hat{Y}$ denotes the predicted outputs. In this work $Y, \hat{Y} \in \mathbb{R}^{T \times 20}$, with ten outputs for each leg. We bootstrapped RRV values 200 times for each output and reported the average RRV values for each output.
